## Supplementary Information for "Identification and characterization of structural and regulatory cell-shape determinants in *Haloferax volcanii*"

**This document includes:**

Extended Data Figures 1-7
Extended Data Tables 1-3
Supplementary Discussions 1 – 7
Legends for Supplementary Data 1 – 3
Legends for Supplementary Videos 1 – 2

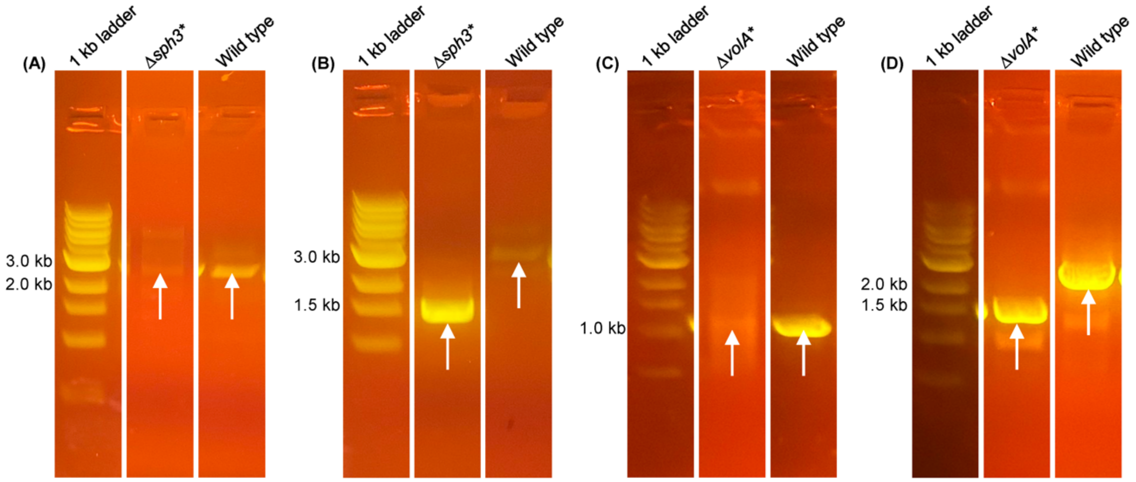

*Extended Data Figure 1: Gel showing partial deletions for* sph3 *and* volA. **(A)** Gel from colony PCR of *sph3* from Δ*sph3** colony using primers 2175_inside_fwd and 2175_KO_down_rvr revealed a faint band at approximately 2.7 kb (1965 bp gene size + ~750 bp downstream of gene), as indicated by white arrows. **(B)** Gel from colony PCR of *sph3* from Δ*sph3** colony using primers 2175_KO_up_fwd and 2175_KO_down_rvr revealed a band at approximately 1.5 kb (~750 bp upstream of gene + ~750 bp downstream of gene) for Δ*sph3** and 3.5 kb (~750 bp upstream of gene + 1965 bp gene size + ~750 bp downstream of gene) for wild type, as indicated by white arrows. **(C)** Gel from PCR of genomic DNA isolated from Δ*volA** using primers 2015_inside_fwd and 2015_inside_rvr revealed a faint band at approximately 1.0 kb (1038 bp gene size), as indicated by white arrows. **(D)** Gel from PCR of genomic DNA isolated from Δ*volA** using primers 2015_KO_up_fwd and 2015_KO_down_rvr revealed a band at approximately 1.5 kb (~750 bp upstream of gene + ~750 bp downstream of gene) for Δ*volA** and 2.5 kb (~750 bp upstream of gene + 1038 bp gene size + ~750 bp downstream of gene) for wild type as indicated by white arrows. All images shown are cut from the original images.

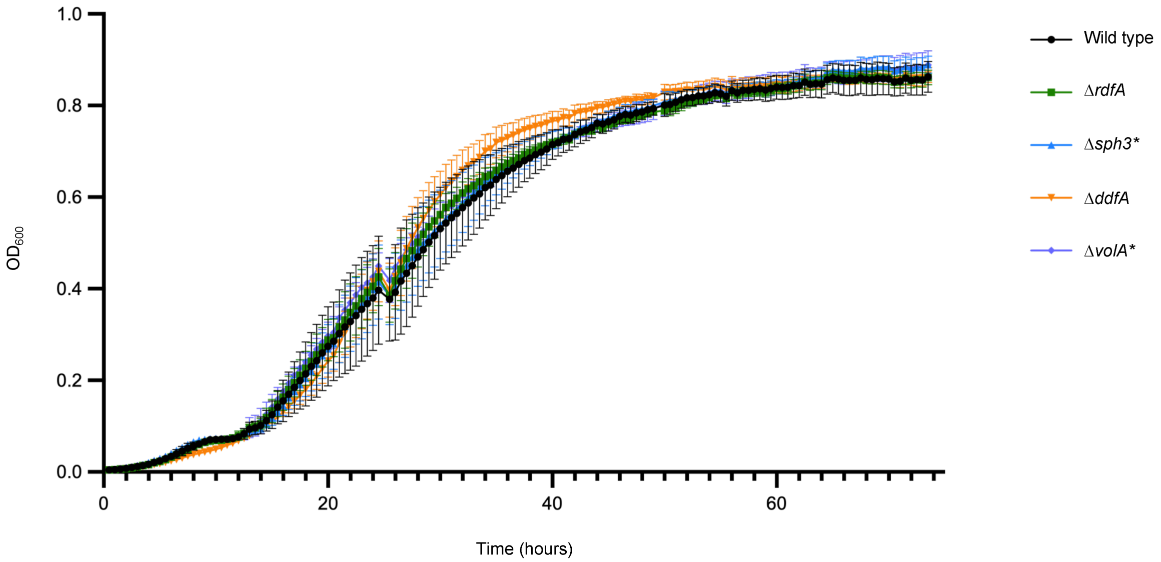

*Extended Data Figure 2: Growth curve for wild type,* ΔrdfA*,* Δsph3**,* ΔddfA*, and* ΔvolA*. Wild-type and mutant strains were grown in a 96-well plate with double orbital shaking for approximately 74 hours, with $OD_{600}$ readings taken every 30 minutes. Error bars are standard deviations across five biological replicates per strain.

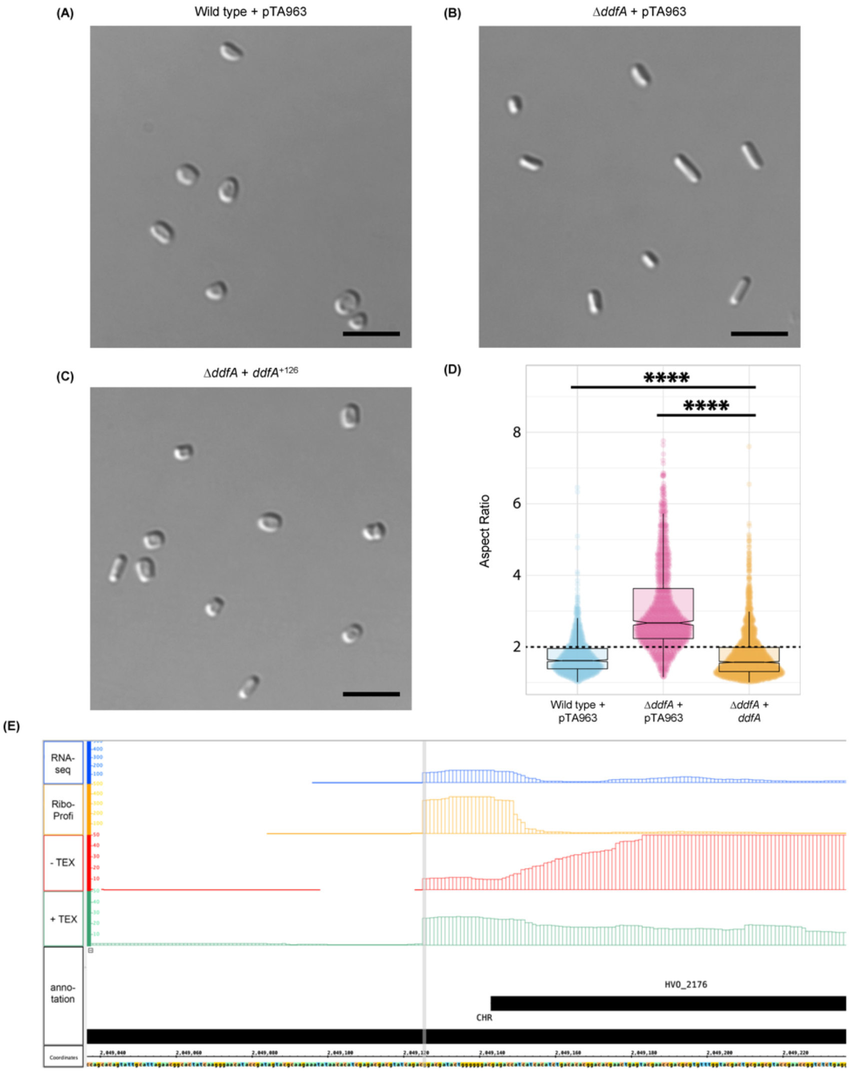

*Extended Data Figure 3: Complementation of* ΔddfA *and predicted start site of* ddfA. Late-log ($OD_{600}$ between 1.7 and 1.8) shape images for **(A)** wild type + empty vector pTA963, **(B)** Δ*ddfA* + empty vector pTA963, and **(C)** Δ*ddfA* + *ddfA*$^{+126}$, each representative of three biological replicates. Δ*ddfA* + *ddfA*$^{+126}$ is the complementation strain; complementation was achieved using the re-annotated *ddfA* gene with an additional 126 nucleotides added to the N-terminus along with a traditional ATG start codon. Scale bars are 5 µm. **(D)** Quantification of cellular aspect ratio. Aspect ratios <2 are considered disks and/or short rods. ****$p<0.0001$. Effect size is -0.041 between wild type + pTA963 and Δ*ddfA* + *ddfA*$^{+126}$ and -1.101 between Δ*ddfA* + pTA963 and Δ*ddfA* + *ddfA*$^{+126}$. **(E)** dRNA-Seq data (1) (green and red panels) reveal a promoter 18 base pairs upstream of the annotated *ddfA* (*hvo_2176*) gene. Red signals (panel -TEX) represent reads from an RNA fraction containing all cellular RNAs. Green signals (panel +TEX) represent reads from an RNA sample treated with terminator 5' phosphate-dependent exonuclease (TEX). Ribosome profiling data (2) are shown in orange (panel Ribo-Profi), and the corresponding RNA-seq data are shown in blue (panel RNA-seq). Ribosome profiling shows an enriched ribosome density in the 5' UTR, typical for a leadered mRNA, which is very likely generated by initiating ribosomes (2). The genome coordinates and the annotation are shown at the bottom in black.

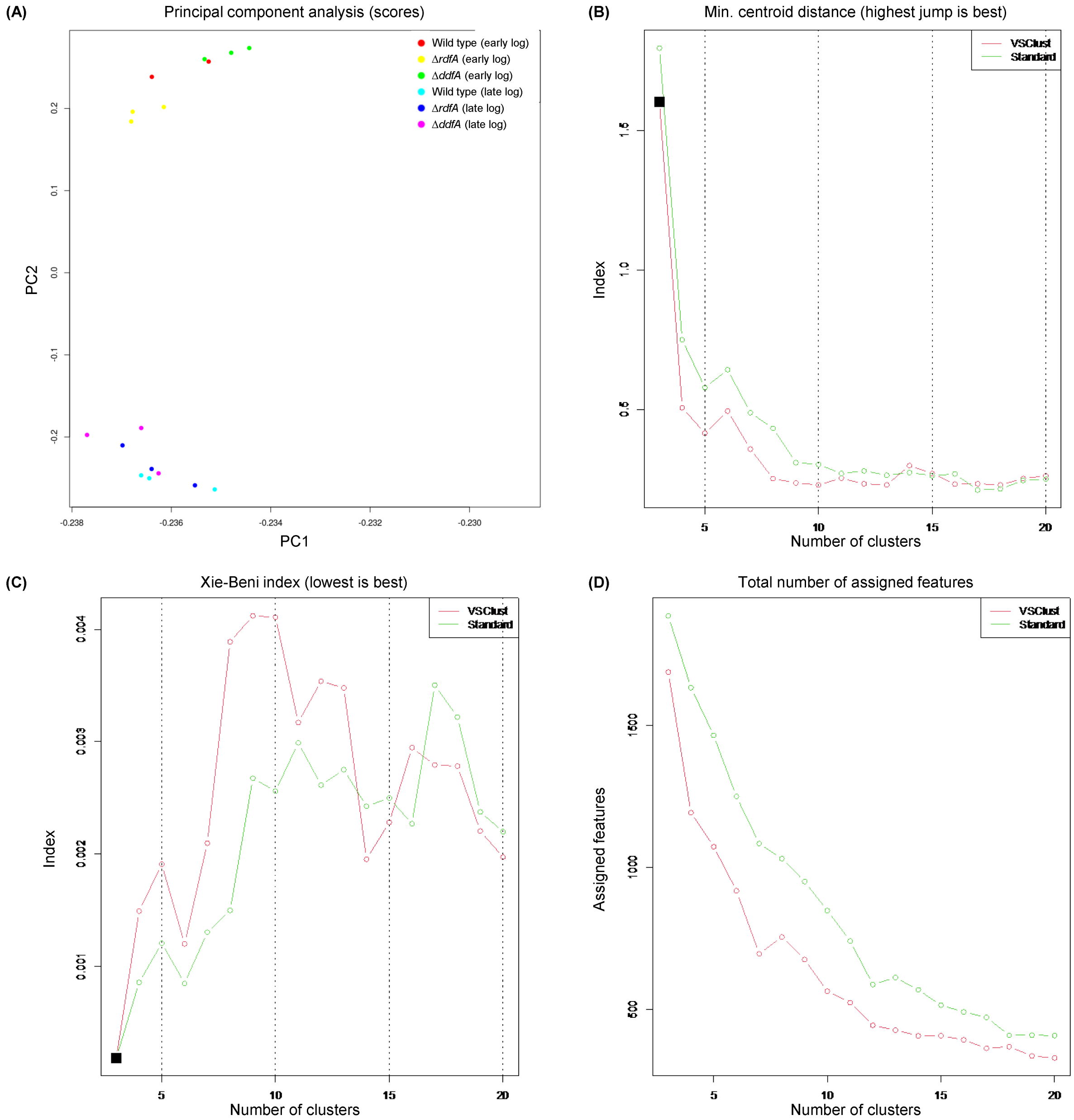

*Extended Data Figure 4: Principal component analysis (PCA) and VSClust estimation of the number of variance-sensitive clusters.* **(A)** PCA plot showing separation of strains and growth phases, with the largest separation observed for PC2, corresponding to differences in the growth phase. Conditions are labeled by color (inserted legend). VSClust reported three metrics for the estimation of cluster numbers: **(B)** Minimum centroid distance, **(C)** Xie-Beni index, and **(D)** total number of assigned features. A cluster number of 14 was chosen because of the corresponding local maximum in (B) and local minimum in (C).

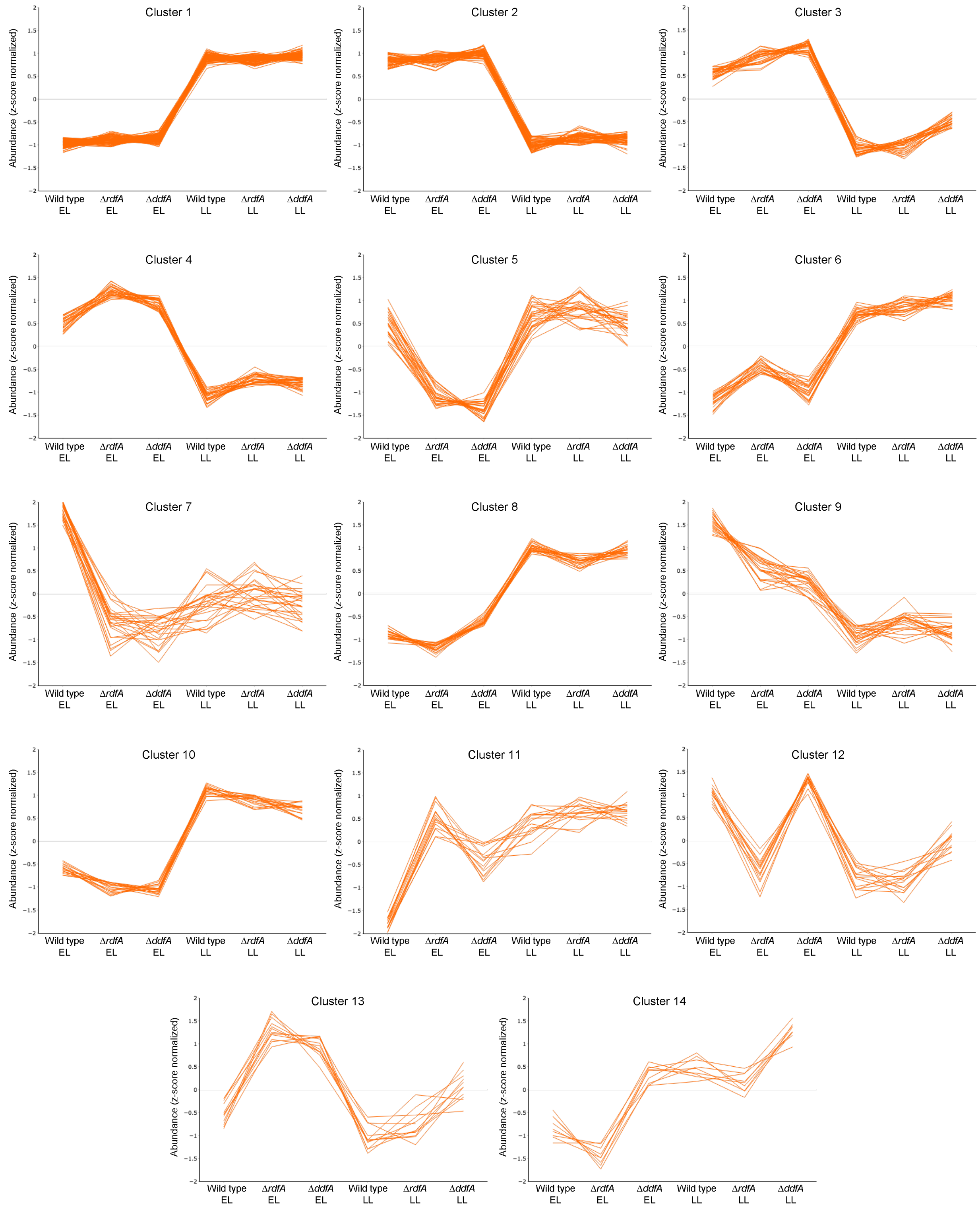

*Extended Data Figure 5: Variance-sensitive clustering based on patterns of protein abundance across strains and growth phases, resulting in 14 clusters.* Normalized protein abundances across each condition are shown for each cluster, with individual proteins or protein groups represented as single lines. 'EL': early-log growth phase, 'LL': late-log growth phase.

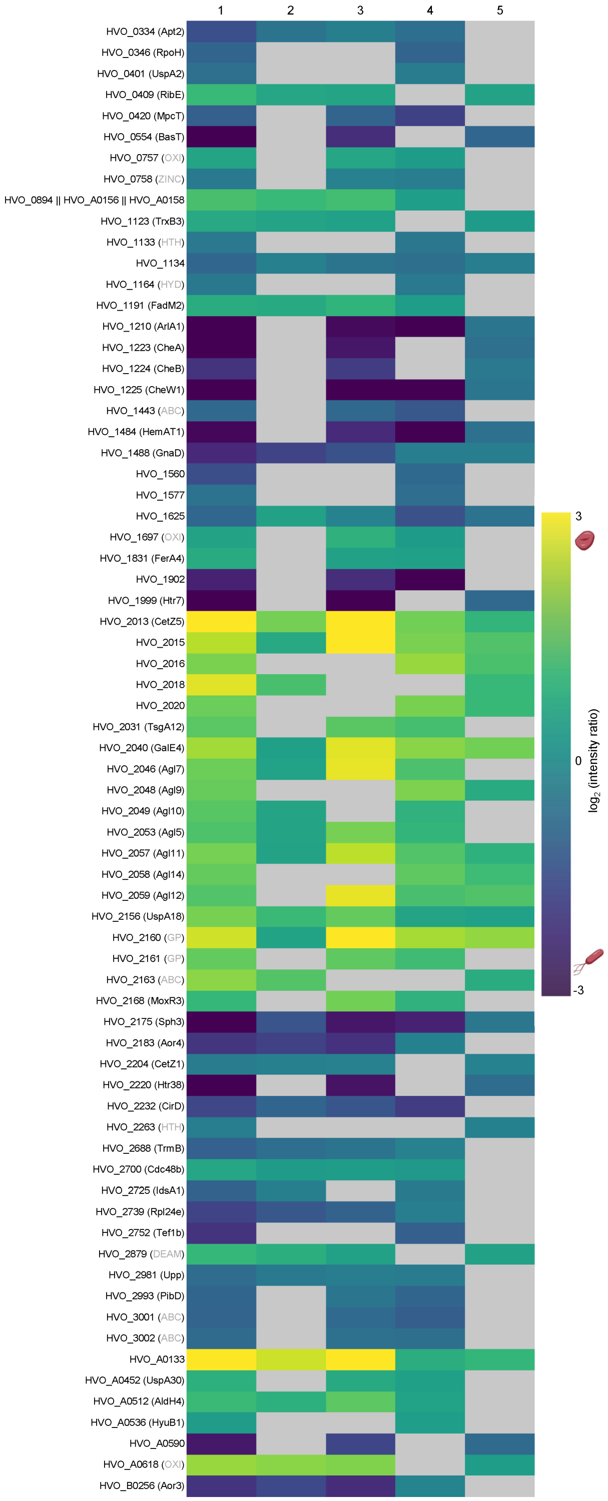

OXI = oxidoreductase; ZINC = zinc-finger protein; HTH = HTH domain-containing protein; HYD = hydrolase; ABC = protein from ABC-type transport system; GP = glycoprotein; DEAM = deaminase

*Extended Data Figure 6: Heatmap of proteins likely important for shape by proteomic comparisons between wild type,* ΔrdfA, *and* ΔddfA. Proteins were filtered for protein abundance ratios with a PEP<0.05 in column 1 and column 4 and/or 5; PEP>0.05 are gray boxes. Column numbers correspond to comparisons defined in Fig. 2A. Protein groups include names for all members, separated by ||.

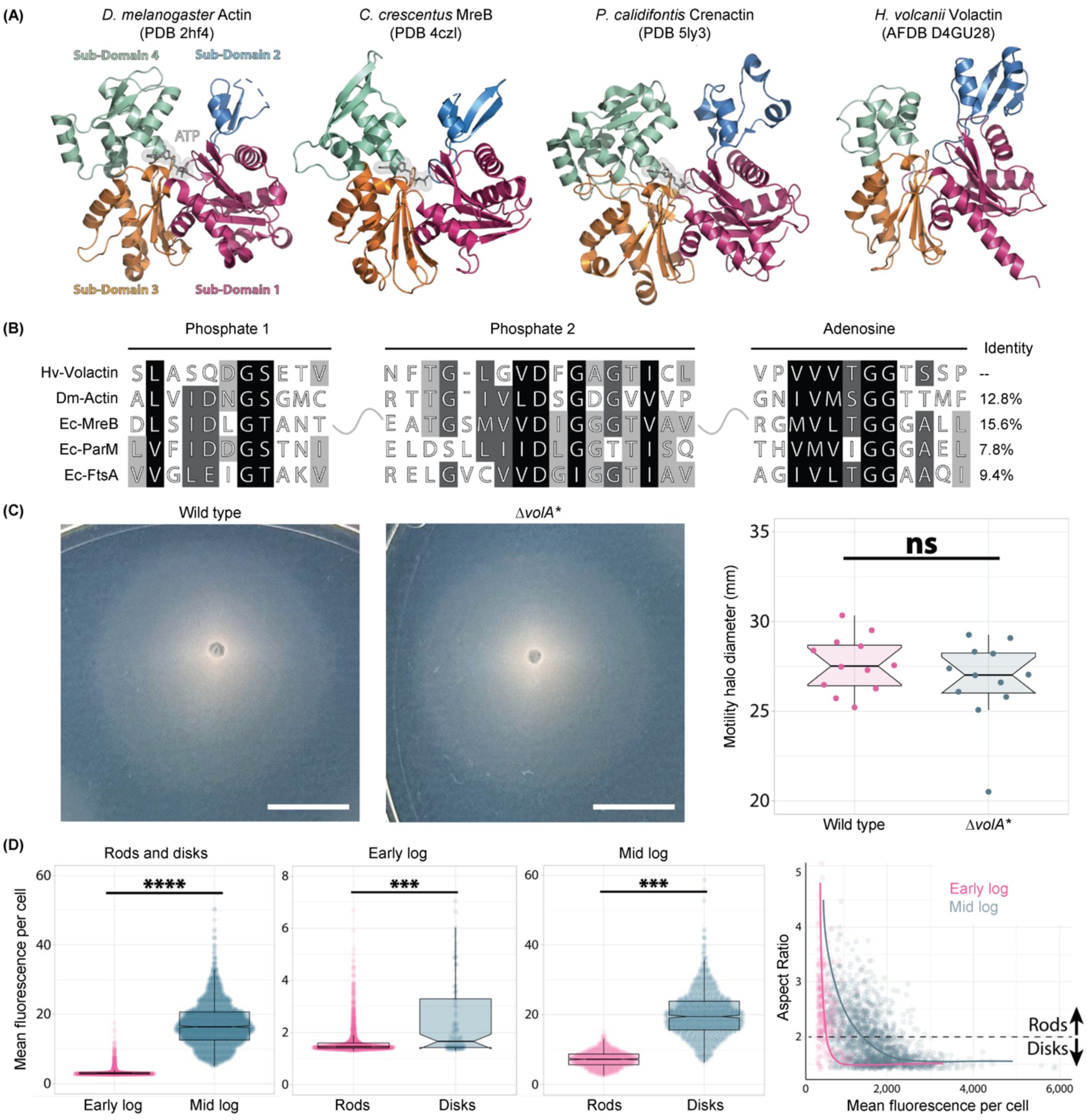

*Extended Data Figure 7: Volactin structural comparison,* ΔvolA* *motility, and quantification of volactin fluorescence in rods and disks between early log and mid log.* **(A)** Structural comparison of actin homologs. From left to right, crystallographic structures of *Drosophila melanogaster* (3), *Caulobacter crescentus* (4), and *Pyrobaculum calidifontis* (5) actins and a structural model for *Hfx. volcanii* volactin as predicted by AlphaFold 2. **(B)** Sequence comparison of nucleotide binding sites of *Hfx. volcanii* volactin, *D. melanogaster* actin, *E. coli* MreB, *E. coli* ParM, and *E. coli* FtsA. **(C)** Left and middle panels, motility halo images for wild type and Δ*volA**. Colonies were stab-inoculated into 0.35% agar and incubated for two days at 45°C and one day at room temperature. Image for each strain is representative of 12 biological replicates. Scale bars indicate 10 mm. Right panel, quantification of motility halo diameter. 'ns' is not significant. **(D)** Plots showing the quantification of GFP-tagged volactin fluorescence in early log versus mid log, early-log rods versus disks, and mid-log rods versus disks as well as the correlation between volactin-msfGFP signal and aspect ratio.

*Extended Data Table 1: Homologs to RdfA and Sph3 (SMC-like protein).* Proteins belong to the same group when they show more than 55% protein sequence identity. Only sequences from the genomes under continuous survey are clustered into groups. In several cases, only the SMC-like protein met the cutoff for grouping. The following organisms are under continuous annotation survey: i) three species from *Haloferax* (HVO, *Hfx. volcanii*; HFX, *Hfx. mediterranei*; HfgLR, *Hfx. gibbonsii*); ii) *Haloquadratum walsbyi* (Hqrw); iii) two species from *Haloarcula* (rrnAC, rrnB and pNG, *Har. marismortui*; HAH, *Har. hispanica*); iv) two species from *Natronomonas* (NP, *Nmn. pharaonis*; Nmlp, *Nmn. moolapensis*); v) *Natrialba magadii* (Nmag); vi) two species from *Halobacterium* (OE, *Hbt. salinarum*; Hhub, *Hbt. hubeiense*); vii) *Halohasta litchfieldiae* (halTADL). The following genomes from 12 additional genera were analyzed: *Halalkalicoccus jeotgali* (HacjB3), *Halogeometricum borinquese* (Hbor), *Halomicrobium mukahataei* (Hmuk), *Halopiger xanaduensis* (Halxa), *Halorhabdus tiamatea* (HTIA), *Halorubrum lacusprofundi* (Hlac), *Haloterrigena turkmenica* (Htur), *Halovivax ruber* (Halru), *Natrinema pellirubrum* (Natpe), *Natronobacterium gregoryi* (Natgr), *Natronococcus occultus* (Natoc), *Salinarchaeum sp.* Harcht-Bsk1 (L593), halophilic archaeon DL31 (Halar).

| Group | Species to which protein belongs | RdfA family | SMC-like protein | Synteny | Comment |
|---|---|---|---|---|---|
| 1 | *Haloferax volcanii* | HVO_2174 (RdfA) | HVO_2175 (Sph3) | adjacent | |
| 2 | *Haloferax volcanii* | HVO_A0179 | HVO_A0180 | adjacent | |
| 3 | *Haloferax volcanii* | HVO_B0117 | HVO_B0118 | adjacent | |
| 4 | *Haloferax volcanii* | HVO_B0174 | HVO_B0173 | adjacent | |
| 1 | *Haloferax mediterranei* | HFX_2233 | HFX_2234 | adjacent | |
| 5 | *Haloferax mediterranei* | HFX_6157 | HFX_6156 | adjacent | |
| 1 | *Haloferax gibbonsii* | HfgLR_12455 | HfgLR_12450 | adjacent | |
| 3 | *Haloferax gibbonsii* | HfgLR_20520 | HfgLR_20525 | adjacent | |
| 6 | *Haloquadratum walsbyi* | Hqrw_1956 | Hqrw_1957 | adjacent | |
| 7 | *Haloquadratum walsbyi* | Hqrw_2741 | - | - | |

| | | | | |
|---|---|---|---|---|
| 6 | *Haloarcula marismortui* | rrnAC2249 | rrnAC2250 | adjacent |
| 8 | *Haloarcula marismortui* | rrnB0193 | - | - |
| 9 | *Haloarcula marismortui* | pNG7224 | pNG7225 | adjacent |
| 10 | *Haloarcula marismortui* | pNG7385 | pNG7386 | adjacent |
| 6 | *Haloarcula hispanica* | HAH_2718 | HAH_2719 | adjacent |
| 8 | *Haloarcula hispanica* | HAH_4309 | - | - |
| 9 | *Haloarcula hispanica* | HAH_5326 | HAH_5325 | adjacent |
| 10 | *Haloarcula hispanica* | HAH_5179 | HAH_5178 | adjacent |
| 11 | *Natronomonas pharaonis* | NP_3408A | NP_3410A | adjacent |
| 12 | *Natronomonas pharaonis* | NP_4260A | NP_4262A | adjacent |
| 12 | *Natronomonas moolapensis* | Nmlp_3023 | Nmlp_3024 | adjacent |
| 13 | *Natrialba magadii* | Nmag_0804 | - | - |
| 14 | *Natrialba magadii* | Nmag_2907 | Nmag_2908 | adjacent |
| 15 | *Halobacterium salinarum* | OE_5048F | OE_5049F | adjacent |
| 16 | *Halobacterium salinarum* | OE_5211F | OE_5212F | adjacent |
| 17 | *Halobacterium hubeiense* | Hhub_2395 | Hhub_2396 | adjacent |
| 18 | *Halobacterium hubeiense* | Hhub_2448 | - | - |

| | | | | | |
|---|---|---|---|---|---|
| 19 | *Halohasta litchfieldiae* | halTADL_0031 | halTADL_0032 | adjacent | |
| 9 | *Halohasta litchfieldiae* | halTADL_2636 | halTADL_2637 | adjacent | |
| 20 | *Halohasta litchfieldiae* | halTADL_2656 | halTADL_2657 | adjacent | |
| 17 | *Halorubrum lacusprofundi* | Hlac_2235 | Hlac_2244 | vicinity | separated by eight genes |
| 18 | *Halorubrum lacusprofundi* | Hlac_2591 | - | - | |
| - | *Halorubrum lacusprofundi* | - | Hlac_2789 | - | |
| 1 | *Halogeometricum boringuese* | Hbor_31810 | Hbor_31800 | adjacent | |
| 3 | *Halogeometricum borinquese* | Hbor_36760 | Hbor_36770 | adjacent | |
| 13 | *Haloterrigena turkmenica* | Htur_0821 | Htur_0820 | adjacent | |
| 9 | *Haloterrigena turkmenica* | Htur_3953 | Htur_3954 | adjacent | |
| - | *Haloterrigena turkmenica* | - | Htur_4125 | - | |
| 12 | *Haloterrigena turkmenica* | Htur_4415 | Htur_4414 | adjacent | |
| - | *Haloterrigena turkmenica* | Htur_4715 | Htur_4714 | adjacent | |
| - | *Halomicrobium mukahataei* | Hmuk_0119 | - | - | |
| - | *Halomicrobium mukahataei* | Hmuk_2049 | Hmuk_2048 | adjacent | |
| - | Halophilic archaeon DL31 | Halar_0328 | Halar_0329 | adjacent | |

| | | | | |
|---|---|---|---|---|
| - | Halophilic archaeon DL31 | Halar_1101 | Halar_1102 | adjacent |
| - | Halophilic archaeon DL31 | Halar_1116 | - | - |
| - | *Halopiger xanaduensis* | Halxa_0987 | - | - |
| - | *Halopiger xanaduensis* | - | Halxa_1877 | - |
| 13 | *Halopiger xanaduensis* | Halxa_2173 | | - |
| 17 | *Natrinema pellirubrum* | Natpe_1605 | Natpe_1604 | adjacent |
| 15 | *Natrinema pellirubrum* | Natpe_1870 | Natpe_1871 | adjacent |
| - | *Natrinema pellirubrum* | Natpe_4419 | Natpe_4420 | adjacent |
| 14 | *Natronobacterium gregoryi* | - | Natgr_1168 | - |
| 13 | *Natronobacterium gregoryi* | Natgr_3161 | - | - |
| 13 | *Natronococcus occultus* | Natoc_0345 | Natoc_0346 | adjacent |
| - | *Natronococcus occultus* | - | Natoc_4085 | - |
| - | *Halalkalicoccus jeotgali* | HacjB3_09675 | HacjB3_09670 | adjacent |
| - | *Halalkalicoccus jeotgali* | - | HacjB3_17161 | - |
| - | *Halalkalicoccus jeotgali* | - | HacjB3_17473 | - |
| - | *Halorhabdus tiamatea* | HTIA_1062 | HTIA_1063 | adjacent |

| | | | | |
|---|---|---|---|---|
| - | *Halorhabdus tiamatea* | HTIA_p2913 | HTIA_p2914 | adjacent |
| - | *Salinarchaeum sp.* Harcht-Bsk1 | L593_06415 | L593_06420 | adjacent |

*Extended Data Table 2: Plasmids, strains, and primers used to construct recombinant plasmids in this study.*

| **Name** | **Relevant characteristic(s)** | **Source** |
|---|---|---|
| Plasmids | | |
| pTA131 | Amp$^r$, pBluescript II with BamHI-XbaI fragments from pGB70 harboring *pfdx-pyrE2* | (6) |
| pTA963 | Amp$^r$, *pyrE2* and *hdrB* markers, Trp-inducible (p.*tna*) promoter | (7) |
| pHS1 | pTA131 carrying fragment with ~1500 nucleotides of fused upstream and downstream regions of *rdfA* | This study |
| pHS2 | pTA131 carrying fragment with ~1500 nucleotides of fused upstream and downstream regions of *sph3* | This study |
| pHS3 | pTA131 carrying fragment with ~1500 nucleotides of fused upstream and downstream regions of *ddfA* | This study |
| pAD12 | pTA131 carrying fragment with ~1500 nucleotides of fused upstream and downstream regions of *hvo_B0194* | This study |
| pAD14 | pTA131 carrying fragment with ~1500 nucleotides of fused upstream and downstream regions of *volA* | This study |
| pHS6 | pTA963 carrying *ddfA* as originally annotated | This study |
| pHS7 | pTA963 carrying extended *ddfA* (*ddfA*$^{+126}$), based on the updated annotation | This study |
| *E. coli* strains | | |
| DH5α | F$^-$ ϕ80Δ*lacZ*ΔM15 (*lacZYA-argF*)*U169 recA1 endA1 hsdR17*($r_K^-$ $m_K^-$) *phoA supE44 thi-1 gyrA96 relA1* | Invitrogen |
| DL739 | MC4100 *recA dam-13*::Tn*9* | (8) |
| *Hfx. volcanii* strains | | |
| H53 | Δ*pyrE2* Δ*trpA* | (6) |
| H98 | Δ*pyrE2* Δ*hdrB* | (6) |
| H295 | H295 (Δ*pyrE2* Δ*trpA* Δ*rad50* Δ*mre11 bga-Ha-Kp*) | (9) |
| H26 | Δ*pyrE2* | (6) |

| | | |
|---|---|---|
| Δ*cetZ1* | H98 Δ*cetZ1* | (10) |
| JK3 | H295 *hvo_B0194::tn* | This study |
| JK5 | H295 *hvo_B0364::tn* | This study |
| SAH1 | H295 *hvo_2176::tn* | (11) |
| HS101 | H53 Δ*rdfA* | This study |
| HS102 | H53 Δ*sph3* (partial) | This study |
| HS103 | H53 Δ*ddfA* | This study |
| AD12 | H53 Δ*hvo_B0194* | This study |
| HS104 | H53 Δ*volA* (partial) | This study |
| HS107 | HS103 containing pHS6 | This study |
| HS108 | HS103 containing pHS7 | This study |
| HS110 | H53 containing empty vector pTA963 | This study |
| HS113 | HS103 containing empty vector pTA963 | This study |
| aBL126 | H26 *volA*::*volA*-40aa-msfGFP-pyrE2 | This study |
| aBL170 | H26 *volA*::*volA*-40aa-msfGFP-pyrE2 ftsZ1::ftsZ1-EG-mApple-I-mevR | This study |
| Primers | 5' to 3' sequence | |
| 2174_KO_up_fwd | attatctagagcttgtggttccgtacgag | This study |
| 2174_KO_up_rvr | gttggtgagcccgaatcgtgtgacttcgccggcg | This study |
| 2174_KO_down_fwd | cgccggcgaagtcacacgattcgggctcaccaac | This study |
| 2174_KO_down_rvr | aatactcgagcgtctaattcggcttggtgc | This study |
| 2175_KO_up_fwd | attattctagatcacgtctgataatccatgggt | This study |
| 2175_KO_up_rvr | gctgtcagggcacgtcaggtgtcggatgtggatgtgtt | This study |
| 2175_KO_down_fwd | aacacatccacatccgacacctgacgtgccctgacagc | This study |
| 2175_KO_down_rvr | aatactcgaggagggcgagcggtcc | This study |
| 2176_KO_up_fwd | attatctagaccacatcgtccgcgcg | This study |

| | | |
|---|---|---|
| 2176_KO_up_rvr | cgaacgacccggcagaggctgcggcgaccgc | This study |
| 2176_KO_down_fwd | gcggtcgccgcagcctctgccgggtcgttcg | This study |
| 2176_KO_down_rvr | attactcgaggagctggatactgctcgttatc | This study |
| B0194_KO_up_fwd | ctagctcgaggtccgcttgatatccgac | This study |
| B0194_KO_up_rvr | tcgcccgtttactggaagcactctggc | This study |
| B0194_KO_down_fwd | tgcttccagtaaacgggcgagtttctg | This study |
| B0194_KO_down_rvr | gatctctagagataagccagtcgaggacg | This study |
| 2015_KO_up_fwd | ctagctcgagccagttcgtcggcgcgtatc | This study |
| 2015_KO_up_rvr | ggaccggcgcacctcacgcgtcgtaggg | This study |
| 2015_KO_down_fwd | cgcgtgaggtgcgccggtccgagctgac | This study |
| 2015_KO_down_rvr | gatctctagacctgccacaccgagtcgaac | This study |
| 2176_OE_fwd | tatattccatatggtggacaaacacgcccgcg | This study |
| 2176_OE_fwd_newannot | tatattcatatgatgggacgatactgggggacg | This study |
| 2176_OE_rvr | tatattggatccttagtgatggtgatggtgatgcgggccgccccggcccgcgtcc | This study |
| 2174_inside_fwd | atgcgagactctggaaggtc | This study |
| 2174_inside_rvr | ctactcgtctcgctcgcg | This study |
| 2175_inside_fwd | atgaacgattcagaggcactt | This study |
| 2176_inside_fwd | gtggacaaacacgcccg | This study |
| 2176_inside_rvr | ttaccggcccgcgtc | This study |
| B0194_inside_fwd | actccgactcaaccgacgca | This study |
| B0194_inside_rvr | gacgaggcggtccgtctcga | This study |
| 2015_inside_fwd | atggcgaaaggccttgacgtagg | This study |
| 2015_inside_rvr | tcagttcgattccggcgcgg | This study |
| oBL174 | gacgaggaagacgaggagg | This study |
| oBL175 | catcctgactcgcgg | This study |
| oBL354 | cttgagggtagcggac | This study |

| | | |
|---|---|---|
| oBL24 | ccatcccctcccatgtcatttgtaaagttcatccattccatg | This study |
| oHV6 | ttagccgtcggcgtc | (12) |
| oHV7 | cgtggataaaacccctcg | (12) |
| oBL232 | cgtagatgcggttgagatg | This study |
| oBL248 | cgaggggttttatccacggcgccggtccgagctga | This study |
| oHV3 | cgtcctccgtaaaccg | This study |
| oHV4 | gtccgctaccctcaagctcgacgtagtcgatgtct | This study |
| oHV140 | gggaggtgacgcctgatggtctcgaagggcg | This study |
| oHV170 | ccatcccctcccatgtcacttgtagagctcgtcc | This study |
| oBL36 | cgaggaagcggaaga | This study |
| oBL37 | catgggaggggatgg | This study |
| oBL308 | ggatctaaatcaaaagaatagaccgatcgagccgtcccg | This study |
| oHV171 | tcttccgcttcctcgtcgagccgtcccg | This study |

*Extended Data Table 3: Parameters for the mass spectrometric analysis of cell shape samples.* A Q Exactive™ HF mass spectrometer (Thermo Fisher Scientific) was employed to measure samples for the comparison between H53, H98, Δ*cetZ1*, and JK3. An Orbitrap Eclipse™ mass spectrometer (Thermo Fisher Scientific) was employed to measure samples for the comparison between H53, Δ*ddfA*, and Δ*rdfA*.

| | Q Exactive™ HF | Orbitrap Eclipse™ |
|---|---|---|
| *Chromatography* | | |
| Column | nanoEase M/Z Peptide BEH C18 column, Waters Corporation, 1.7 um particle size, 75 um x 250 mm | |
| Column oven | 50°C | |
| Flow rate | 300 nl/min | |
| Buffer system | Buffer A: 0.1% formic acid in $H_2O$<br>Buffer B: 0.1% formic acid in acetonitrile | |
| Gradient | 1 min 3% B;<br>increase to 17% B over 39 min;<br>increase to 32% B over 20 min;<br>increase to 90% B over 2 min;<br>9 min 90% B;<br>decrease to 3% B over 2 min;<br>17 min 34% B | |
| *Mass spectrometry, general settings* | | |
| Ion mode | positive | |
| Excluded charge states | 1, >6 | |
| Dynamic exclusion | 10 s | |
| *MS1 settings* | | |
| Resolution | 60,000 | 120,000 |
| Maximum injection time | 100 ms | 50 ms |
| AGC target | 1e6 | 250% |

| | | |
|---|---|---|
| Detector | Orbitrap | |
| Scan range | 400-2000 *m/z* | |
| *MS2 settings* | | |
| Resolution | 15,000 | |
| Detector | Orbitrap | |
| AGC target | 1e5 | 200% |
| Maximum injection time | 200 ms | dynamic |
| Data-dependent acquisition | TopN with N=20 | cycle time 3 s |
| Scan range | fixed first 130 *m/z* | automatic |
| Fragmentation | Higher-energy collisional dissociation | |
| Normalized collision energy | stepped: 25, 30, 35 | |

*Supplementary Discussion 1: Limitations of using genetic screens for identification of shape- and motility-related genes.*

Transposon mutant screens for hypermotile mutants allowed for the isolation of mutant strains with shape defects, thus enabling identification of genes potentially important for both motility and shape. However, results from these screens are inherently limited to genes that affect both motility and shape and are non-essential, having, at most, minor effects on growth rates. These limitations of genetic screens prevent their use for more comprehensive analyses of shape transition pathways and, together with their laborious nature, render genetic screens less suitable for a more comprehensive identification of proteins involved in cell-shape pathways.

*Supplementary Discussion 2: Strain descriptions.*

Strain H53 (Δ*pyrE2* Δ*trpA*) is used as the parent strain for all clean gene deletions and is referred to as "wild type" throughout this manuscript. Strain H53 is derived from strain H26 (Δ*pyrE2*) (6). The Δ*cetZ1* mutant is based on parent strain H98 (Δ*pyrE2* Δ*hdrB*), which is also derived from strain H26 (6). The additional selection marker *hdrB* is expected to be neutral with respect to cell shape.

Strains JK3, JK5, and SAH1 are mutants of the transposon insertion library (13). The background of this library is strain H295 (Δ*pyrE2* Δ*trpA* Δ*rad50* Δ*mre11 bga-Ha-Kp*) (9). The genealogy of strain H295 is strain H204 (Δ*pyrE2* Δ*rad50* Δ*mre11 bga-Ha-Kp*), strain H115 (Δ*pyrE2 bga-Ha-Kp*), strain H54 (Δ*pyrE2 bga-Ha*), strain H26 (9).

It should be noted that the Δ*trpA* mutation of strain H295 was complemented during generation of the transposon insertion library, as the transposon constructs carry the *trpA* gene as a selection marker. The replacement of the *bgaH* gene of *Hfx. volcanii* by a mutant version of the gene from *Haloferax alicantei* was made to allow for assays related to *rad50* and *mre11* (9). It is highly unlikely that this mutation has any shape effect. Usage of a strain mutated in Δ*rad50* and Δ*mre11* had been considered helpful for generation of a transposon insertion library. These mutations are also likely to be neutral with respect to cell shape.

Thus, even though distinct parents have been used for the mutants, all of them differ by only a few genes, as they are all derived from the same ancestral strain H26 (Δ*pyrE2*) (6). The genealogy of strain H26 has been recently described (11). Strain H26 is based on strain DS70 (14), which is a direct descendent of wild-type strain $DS2^T$ (15, 16). To generate strain DS70, the wild-type strain was cured of the small plasmid pHV2 (14). During curing, plasmid pHV4 has been inadvertently integrated into the chromosome (17).

*Supplementary Discussion 3: Comparative quantitative proteomics analysis of strains JK3,* ΔcetZ1*, and wild type.*

Quantitative proteomics was performed using wild-type strains H53 and H98 and shape-defective strains JK3 and Δ*cetZ1*. After identification and label-free quantification of 1944 proteins, a differential abundance in at least one comparison could be found for 314 proteins (Supplementary Data 1). For most of these proteins (n=218), significant changes in abundance were only seen for comparisons within a single strain across growth phases. However, 46 proteins showed differential abundances for comparisons between early- and late-log wild-type cells as well as at least one shape-specific comparison between wild-type and shape-mutant cells within a single growth phase (Supplementary Data 1). These proteins were further considered as potential candidates for proteins with a shape-related function.

*Supplementary Discussion 4: The syntenic association of RdfA and Sph3 homologs is highly conserved.*

SyntTax analysis of RdfA uncovered a strong syntenic coupling to Sph3. RdfA defines a protein family (RdfA family) with a total of four paralogs in *Hfx. volcanii* (Extended Data Table 1). Analyzing 24 genomes from 19 haloarchaeal genera, 54 RdfA family proteins and 50 SMC-like proteins were identified. The majority of the RdfA family proteins (42 of 54, 78%) and the SMC-like proteins (42 of 50, 84%) are encoded directly adjacent to each other. There are only 12 RdfA family proteins and eight SMC-like proteins that lack a partner gene. In one of these cases, the two proteins are encoded in genomic vicinity with an intercalation of eight unrelated genes.

UniProt was used to look up the 54 RdfA family proteins, but InterPro domain assignments were not encountered, with a single exception (NP_3408A), which likely carries a false positive domain assignment. RdfA itself is named 'ASCH domain-containing protein' in UniProt but lacks an assignment of that domain by InterPro. In the current UniProt release (accessed Feb-2023), all proteins receive some name assignment by an artificial intelligence system (UnProtein). Although all members of the RdfA family are related to each other and thus are speculated to fulfill the same or a similar function, a total of 27 distinct protein names have been assigned in UniProt for the 54 members analyzed (e.g. HalOD2 domain-containing protein; Integrase; Minichromosome maintenance protein MCM; peptidyl-prolyl cis-trans isomerase; Portal protein; Tail assembly chaperone; Transcription initiation factor TFIIE). We propose that homologs of RdfA which have not been subjected to experimental analysis are annotated as ‘RdfA family protein’, and only experimentally characterized members are given more specific names (e.g. rod-determining factor RdfA).

*Supplementary Discussion 5: Re-annotation of DdfA.*

An attempt to complement Δ*ddfA* with a copy of *ddfA* carried on a plasmid was unsuccessful, leading us to characterize its gene annotation more carefully. Unlike most haloarchaeal DdfA homologs, the annotated *Hfx. volcanii* protein lacks approximately 126 nucleotides at the N-terminus. Notably, this annotation amendment places the SAH1 transposon insertion within rather than upstream of *ddfA*. Hence, we expressed DdfA with the traditional start codon ATG inserted upstream of its first attributable codon, GGA (*ddfA*$^{+126}$). Cells expressing the plasmid with *ddfA*$^{+126}$ formed disks, confirming that DdfA is required for disk formation (Extended Data Fig. 3a-c). Subsequent transcriptional analysis revealed that the *ddfA* transcription start site may actually be located 18 nucleotides upstream of the annotated start site, but there is no traditional start codon in the vicinity (Extended Data Fig. 3e). According to the Ribosome profiling data, the start codon could be GGG, located six nucleotides upstream of the annotation start. However, the in vivo start codon of DdfA has yet to be identified.

*Supplementary Discussion 6: Comparative quantitative proteomics analysis of strains* ΔrdfA*,* ΔddfA*, and wild type.*

The proteomic analysis of JK3 and Δ*cetZ1*, and subsequent reverse genetics approach, provided a proof-of-concept for the identification of proteins that are important for rods and disks. RdfA and DdfA were identified and confirmed in their roles in cell-shape transition, and deletion mutants for their encoding genes were used for subsequent proteomics. This in-depth proteomic analysis resulted in the identification and label-free quantification of 2328 proteins, with 938 proteins showing differential abundances in at least one comparison (Supplementary Data 3). Knowledge-based filtering to identify proteins of interest, specifically ones that showed differential abundances in comparisons for wild-type cells across growth phases along with at least one comparison between a shape mutant and wild type, resulted in 71 proteins (Supplementary Data 3). 50 of these were not identified in this category in the initial proof-of-concept analysis, illustrating the benefits of using Δ*rdfA* and Δ*ddfA* mutants and the optimized mass spectrometry approach.

In addition to knowledge-driven comparisons based on the phenotypes of the different strains, we also performed explorative data processing using principal component analysis (PCA) and variance-sensitive clustering based on normalized abundances for each condition and strain. The PCA showed a strong separation of samples from early log and late log as well as between wild type and Δ*rdfA* samples from early log and wild type and Δ*ddfA* samples from late log, which is in line with observed phenotypes (Extended Data Fig. 4a). However, the separation between early- and late-log samples for all strains was substantially more pronounced than the differences between strains within a growth phase, indicating that the proteomes of the shape mutants were still strongly altered in different growth phases.

The variance-sensitive clustering analysis (Extended Data Fig. 4c-d; Extended Data Fig. 5; Supplementary Data 2) showed protein abundance patterns for each condition and strain and thus enabled identification of specific clusters with proteins that may serve similar functions or may be part of the same pathway. For six of the fourteen clusters, differences in abundance were mostly due to changes in growth phase, with only minor differences between the strains. They could be grouped into clusters of proteins with higher abundance in late log (clusters 1, 8, and 10; Extended Data Fig. 5) or higher abundance in early log (cluster 2, 3, and 4; Extended Data Fig. 5). Proteins in these clusters were likely involved in cellular processes that correspond to the respective growth phase but were unlikely to be important for shape.

In contrast, the protein abundance patterns of clusters 11, 12, and 14 indicated that the corresponding proteins were involved in cell-shape determination (see main results). Interestingly, for cluster 11, protein abundances in late log were comparable between all strains, at a level similar to Δ*rdfA* in early log (Fig. 4a). Given that some Δ*ddfA* rods are shortened in late log, perhaps cells lacking *ddfA* initiate the transition to disks by forming short, rounded rods in late log, but DdfA is required to complete the transition to disks. We therefore hypothesize that cluster 11 comprises proteins that are involved in the early stages of disk transition and that DdfA is crucial for a later step in the pathway to form disks.

While cluster 6 showed an overall similar abundance pattern as cluster 11, it was less pronounced, with only slightly higher abundances in early log for proteins from Δ*rdfA* compared to wild type and Δ*ddfA* (Extended Data Fig. 5). The roles of these proteins in disk-shaped cells therefore remain to be elucidated. The protein abundance patterns for the remaining four clusters (5, 7, 9, and 13) showed differences for both shape mutants compared to wild type (Extended Data Fig. 5). Since both mutants have distinct shapes, it is unlikely that these clusters correspond to proteins involved in shape-specific processes. Instead, they might be part of stress responses as a result of their altered shapes.

The proteins present in shape-specific clusters revealed potential connections between shape and adhesion. Two proteins in cluster 11, GdhA1 (HVO_1451), a glutamate dehydrogenase, and hypothetical protein HVO_2447, have been shown before to likely be involved in surface adhesion, as transposon insertions in the genes of either protein result in adhesion defects (18). Thus, the regulation of cell-shape transitions may be important for cellular adhesion to surfaces. Moreover, zinc-finger protein HVO_0758 was present in cluster 12, and previous work has shown that deletion of *hvo_0758* results in a loss in swarming ability along with an increase in biofilm formation relative to wild type (19), the latter of which aligns with the fact that genes potentially important for disk formation result in a defect in adhesion when absent.

Two additional proteins that are likely important for shape are SMC-like protein Sph4 (HVO_B0173) and hypothetical protein HVO_1134. Sph4 was a member of cluster 14, indicating its involvement in genetic regulation for the transition to disks. We show that Sph3 is important for shape, and thus, more broadly, SMC-like proteins in general may have shape effects. HVO_1134 was a member of cluster 12 and thus is likely involved in rod formation. HVO_1134 was also indicated to be important for rods based on the knowledge-based comparisons of wild type and shape mutants. While its exact role remains to be elucidated, it is annotated as a transcriptional regulator by UnProtein.

It is worth noting that the clustering was based on protein abundance patterns across all strains and conditions, and relatively small differences in these patterns, even for a single condition, can result in the exclusion of a protein from a cluster. In contrast, the knowledge-based filtering for proteins that showed differential abundances in shape-specific comparisons (Extended

Data Fig. 6; Supplementary Data 1 and 3) can result in proteins with a variety of abundance patterns. Therefore, together, the clustering and knowledge-based filtering provide a more comprehensive picture of proteins that are likely involved in shape transitions.

For example, the transducers MpcT (HVO_0420) and Htr7 (HVO_1999), which showed higher abundances in rods relative to disks in comparisons between wild type and Δ*rdfA* (Extended Data Fig. 6), did not reach the significance threshold to be considered part of a cluster (Supplementary Data 2). However, their abundance pattern was most similar to that of proteins in cluster 12 (Fig. 4b), indicating that these proteins might be involved in signaling cascades that are not just related to chemotaxis but also to shape transitions. Similarly, Sph3 and CetZ1 were only weakly associated with cluster 12 but showed significant abundance differences in shape-specific comparisons (Extended Data Fig. 6), and the phenotypes of their respective mutants Δ*sph3** and Δ*cetZ1* (10) confirmed their roles in shape determination.

In addition, various ABC transporters showed differential abundances in at least one shape-specific comparison (Extended Data Fig. 6). Two of them, HVO_1442 and HVO_3002, were part of cluster 12, and their respective adjacent genes HVO_1443 and HVO_3001 showed similar abundance patterns (Supplementary Data 2). However, the ABC transporter HVO_1705 was part of cluster 4 (Supplementary Data 2), indicating that its function might be related to changes in the growth phase rather than shape. Other ABC transporters (HVO_2031, HVO_2163) were not clearly associated with a specific cluster (Supplementary Data 2). The substrates for any of these transporters remain to be elucidated.

Further analysis of the proteins that showed a likely importance in shape determination based on the knowledge-based filtering using comparisons between wild type and shape mutants (Extended Data Fig. 6; Supplementary Data 1 and 3) revealed several genomic regions that encode for multiple proteins likely involved in shape determination. The importance of the genomic region between *hvo_2160* and *hvo_2176* (*ddfA*), was indicated by the performed genetic screens as well as the proteomic comparisons including JK3 and Δ*cetZ1*, and additional genes were suggested through the proteomic analysis of Δ*rdfA* and Δ*ddfA*. The glycoproteins HVO_2160 and HVO_2161, the ABC transporter HVO_2163, and the potential AAA-type ATPase MoxR (HVO_2168), showed higher abundances in disks, while RdfA (HVO_2174) and Sph3 (HVO_2175) were more abundant in rods. While no quantitative proteomics data could be obtained for DdfA, potentially due its small size, its deletion resulted in a rod-only phenotype (see main results).

Another genomic region encoding for proteins with potential roles in cell shape is that which covers various enzymes of the Agl15-dependent *N*-glycosylation pathway: Agl5, Agl7, Agl9, Agl10, Alg11, Agl12, and Agl14. All of these were significantly more abundant in disks than rods (Extended Data Fig. 6). While none of them was part of a cluster, their abundance patterns were similar to clusters 6 and 11 (Supplementary Data 2), suggesting importance for disk formation. Two *N*-glycosylation pathways exist in *Hfx. volcanii*: AglB-dependent and Agl15-dependent *N*-glycosylation, and deletion mutants of *aglB* or *agl15* have been shown to have shape defects (20). Specifically, while Δ*agl15* makes rods longer into mid- and late-log growth phases than wild type, suggesting the importance of the Agl15-dependent *N*-glycosylation pathway in disk formation, Δ*aglB* makes more disks in early log than wild type, implicating the AglB-dependent *N*-glycosylation pathway in rod formation (20). Our results, in combination with previous work, suggest that different shapes may be associated with distinct *N*-glycans. Specifically, Agl15-dependent *N*-glycosylation seems to be involved in the transition to disks, but its exact role remains to be elucidated. It is possible that specific *N*-glycans allow for cellular processes to occur that are specific to certain shapes, such as motility, adhesion, or nutrient acquisition. However, it is important to note that *N*-glycosylation pathways might not be direct effectors of cell-shape transition; rather, any changes in type or extent of *N*-glycosylation might be an adaptation which occurs after the transition and thus may be downstream of the actual shape change. In a separate analysis, we identified and quantified multiple *N*-glycoproteins. However, the significant protein abundance differences (PEP<0.05) in shape-relevant comparisons were only observed for *N*-glycosylated ArlA1, ArlA2, as well as HVO_2160. The abundance ratios for the corresponding *N*-glycopeptides were similar to the non-glycosylated peptides of the same protein; thus, a specific effect on the degree of *N*-glycosylation could not be concluded. It should be noted that all reliably quantified *N*-glycopeptides were of the AglB-dependent type, and enzymes of the AglB-dependent glycosylation pathway were not observed in shape-related clusters or in lists of candidate proteins after knowledge-driven filtering. Conversely, enzymes of the Agl15-dependent *N*-glycosylation pathway were strong candidates to be involved in shape, but only a handful of *N*-glycopeptides with Agl15-dependent modifications were identified to date (20), and none of them could be reliably quantified here.

Lastly, the third region between *hvo_2013* and *hvo_2020* encodes for proteins CetZ5 (HVO_2013) and VolA (HVO_2015) along with uncharacterized proteins HVO_2016, HVO_2018, and HVO_2020, all of which showed higher abundance in disks (Extended Data Fig. 6). This genomic region was particularly interesting, since VolA was also part of the five proteins that showed significant abundance differences across all shape-specific comparisons, not only for those including Δ*rdfA* and Δ*ddfA* but also for JK3 and Δ*cetZ1.*

Surprisingly, the results of the quantitative proteomics using Δ*rdfA* and Δ*ddfA* did not indicate an importance for shape for HVO_B0194, the protein whose gene was disrupted by a transposon insertion in the mutant strain JK3. Subsequent generation of a deletion strain Δ*hvo_B0194* revealed that it did not have a shape defect (data not shown), supporting the proteomics results and suggesting that other, secondary genome alterations in JK3 were responsible for the rod-only phenotype.

*Supplementary Discussion 7: Identification and characterization of volactin.*

Volactin was originally identified by BLAST searching the *Hfx. volcanii* genome with annotated actin homolog sequences from various archaeal and bacterial species. The *Halapricum salinum* FtsA-like protein, WP_049993655.1, retrieved WP_004043705.1 entered as "hypothetical protein" in the *Hfx. volcanii* genome. Using SwissModel to fit the sequence to the crystallographic structure of MreB from *Thermotoga maritima* confirmed that the *Hfx. volcanii* sequence was compatible with an actin-like fold.

To further investigate HVO_2015 (volactin), as it was selected as a strong candidate in both rounds of proteomics, we explored its predicted AlphaFold structure (21) from the EMBL-EBI Database (https://alphafold.ebi.ac.uk/entry/D4GU28). Using the Dali server (22) to seek for possible homologs, we obtained a vast majority of actin-related proteins, with the top hit being the bacterial rod shape-determining MreB (23). This finding is in line with results from our previous BLAST search. We compared the predicted structure of HVO_2015 with experimentally solved actin structures across eukaryotic and prokaryotic species (Extended Data Fig. 7a), revealing strong similarities in their domain organization. Additionally, HVO_2015 sequence alignment with eukaryotic and bacterial actin families showed conserved ATP-binding motifs that were previously characterized (24), specifically Phosphate 1 (xhxh**D/E**xGpxx), Phosphate 2 (xssxhhh**D**x**G**ssshxx), and Adenosine (xxhhhx**GG**ssxx) (Extended Data Fig. 7b).

Analyzing images of cells expressing volactin-msfGFP at early- (primarily rods) and mid-log (mixed rods and disks) growth phases, we observed volactin filaments in both populations (Fig. 5e). However, a higher fluorescence signal from volactin-msfGFP polymers was observed in mid-log (μ=855±322) cells compared to early-log (μ=169±65) cells (Extended Data Fig. 7d). To determine whether this difference is related to growth phase or disk enrichment in the mid-log population, we used machine learning-based image segmentation and cellular aspect ratio to distinguish rods from disks (12). Supporting our proteomics data, we concluded that disks exhibited approximately a 2-fold increase in volactin-msfGFP signal compared to rods in early-log ($\mu$=340±161 and $\mu$=166±58, respectively) and a 3-fold increase in mid-log ($\mu$=2023±643 and $\mu$=764±314, respectively) growth phases (Extended Data Fig. 7d). Interestingly, plotting the correlation between volactin-msfGFP signal and aspect ratio showed a skewed sub-population of rods with more volactin polymer material at mid- compared to early-log growth phase (Extended Data Fig. 7d).

As described in the main text, volactin was found to be involved in the timely transition to disks. However, the exact mechanism of volactin function in the cell has yet to be elucidated. Volactin is not the only factor known to be involved in the timely disk transition. Depletion of enzymes such as archaeaosortase A (ArtA), phosphatidylserine synthase (PssA), decaboxylase (PssD), and the protease LonB all result in a larger proportion of rod-shaped cells than wild type in mid log (12, 25). Although ArtA, PssA, and PssD were shown to localize with the cytokinetic ring (12), future studies should explore whether volactin filaments are controlling proteolysis and development in space and time. Likewise, it will be important to understand whether substrates are transported along the volactin filaments, which span the cytoplasm from one inner membrane region to another, and if so, what these substrates are.

Concomitantly with our work, salactin, another actin filament system, was characterized in the haloarchaeon *Hbt. salinarum* (Zheng et al., in preparation). Salactin polymers are dynamically unstable, similar to our observations for volactin (Fig. 5c; Supplementary Video 1), but salactin filaments are anchored at the cell poles and polymerize towards the cytoplasm. Intriguingly, unlike volactin, salactin plays no role in cell morphogenesis but rather aids in DNA partitioning when chromosomal copy number is low. The identification of these two functionally distinct classes of halophilic actins suggests that additional archaeal cytoskeletal proteins may be dedicated to very specific cellular processes.

*Supplementary Data 1: Proteins identified through proteomic comparisons with wild type, JK3, and* ΔcetZ1. File includes all quantified proteins (tab 1), proteins that have significant abundance differences for at least one comparison (tab 2), and proteins that have significant abundance differences for wild type between early log and late log as well as at least one shape-specific comparison (tab 3).

*Supplementary Data 2: Proteins identified and sorted into clusters.* List of proteins classified within each of the 14 variance-sensitive clusters.

*Supplementary Data 3: Proteins identified through proteomic comparisons with wild type,* ΔrdfA*, and* ΔddfA. File includes all quantified proteins (tab 1), proteins that have significant abundance differences for at least one comparison (tab 2), proteins that have significant abundance differences for wild type between early log and late log as well as at least one shape-specific comparison (tab 3), and proteins that have significant abundance differences for wild type between early log and late log as well as both shape-specific comparisons (tab 4). Proteins that show significant abundance differences for wild type between early log and late log as well as both shape-specific comparisons in both proteomics experiments (JK3, Δ*cetZ1*, and wild type along with Δ*rdfA*, Δ*ddfA*, and wild type) are listed in tab 5.

*Supplementary Video 1: Volactin filament assembly and disassembly is independent of the division site.* Colocalization of volactin- and FtsZ-tagged protein fluorescence reveals a lack of overlap between the two proteins, indicating spatial separation between volactin filament assembly and disassembly and cell division.

*Supplementary Video 2: Volactin filaments bind the membrane.* 3-D super-resolution projections reveal volactin filaments span the cytoplasm and are bound from their tips to the membrane.

References

1. Babski J, Haas KA, Näther-Schindler D, Pfeiffer F, Förstner KU, Hammelmann M, Hilker R, Becker A, Sharma CM, Marchfelder A, Soppa J. 2016. Genome-wide identification of transcriptional start sites in the haloarchaeon Haloferax volcanii based on differential RNA-Seq (dRNA-Seq). BMC Genomics 17:629.

2. Hadjeras L, Bartel J, Maier L-K, Maaß S, Vogel V, Svensson SL, Eggenhofer F, Gelhausen R, Müller T, Alkhnbashi OS, Backofen R, Becher D, Sharma CM, Marchfelder A. 2023. Revealing the small proteome of *Haloferax volcanii* by combining ribosome profiling and small-protein optimized mass spectrometry. microLife 4:uqad001.

3. Rould MA, Wan Q, Joel PB, Lowey S, Trybus KM. 2006. Crystal Structures of Expressed Non-polymerizable Monomeric Actin in the ADP and ATP States. J Biol Chem 281:31909–31919.

4. van den Ent F, Izoré T, Bharat TA, Johnson CM, Löwe J. 2014. Bacterial actin MreB forms antiparallel double filaments. eLife 3:e02634.

5. Izoré T, Kureisaite-Ciziene D, McLaughlin SH, Löwe J. 2016. Crenactin forms actin-like double helical filaments regulated by arcadin-2. eLife 5:e21600.

6. Allers T, Ngo H-P, Mevarech M, Lloyd RG. 2004. Development of Additional Selectable Markers for the Halophilic Archaeon Haloferax volcanii Based on the leuB and trpA Genes. Appl Environ Microbiol 70:943–953.

7. Allers T, Barak S, Liddell S, Wardell K, Mevarech M. 2010. Improved Strains and Plasmid Vectors for Conditional Overexpression of His-Tagged Proteins in Haloferax volcanii. Appl Environ Microbiol 76:1759–1769.

8. Blyn LB, Braaten BA, Low DA. 1990. Regulation of pap pilin phase variation by a mechanism involving differential dam methylation states. EMBO J 9:4045–4054.

9. Delmas S, Shunburne L, Ngo H-P, Allers T. 2009. Mre11-Rad50 Promotes Rapid Repair of DNA Damage in the Polyploid Archaeon Haloferax volcanii by Restraining Homologous Recombination. PLoS Genet 5:e1000552.

10. Duggin IG, Aylett CHS, Walsh JC, Michie KA, Wang Q, Turnbull L, Dawson EM, Harry EJ, Whitchurch CB, Amos LA, Löwe J. 2015. CetZ tubulin-like proteins control archaeal cell shape. Nature 519:362–365.

11. Collins M, Afolayan S, Igiraneza AB, Schiller H, Krespan E, Beiting DP, Dyall-Smith M, Pfeiffer F, Pohlschroder M. 2020. Mutations Affecting HVO_1357 or HVO_2248 Cause Hypermotility in Haloferax volcanii, Suggesting Roles in Motility Regulation. Genes 12:58.

12. Abdul-Halim MF, Schulze S, DiLucido A, Pfeiffer F, Filho AWB, Pohlschroder M. 2020. Lipid Anchoring of Archaeosortase Substrates and Midcell Growth in Haloarchaea 11:14.

13. Kiljunen S, Pajunen MI, Dilks K, Storf S, Pohlschroder M, Savilahti H. 2014. Generation of comprehensive transposon insertion mutant library for the model archaeon, Haloferax volcanii, and its use for gene discovery. BMC Biol 12:103.

14. Wendoloski D, Ferrer C, Dyall-Smith ML. 2001. A new simvastatin (mevinolin)-resistance marker from Haloarcula hispanica and a new Haloferax volcanii strain cured of plasmid pHV2. Microbiology 147:959–964.

15. Mullakhanbhai MF, Larsen H. 1975. Halobacterium volcanii spec. nov., a Dead Sea halobacterium with a moderate salt requirement. Arch Microbiol 104:207–214.

16. Hartman AL, Norais C, Badger JH, Delmas S, Haldenby S, Madupu R, Robinson J, Khouri H, Ren Q, Lowe TM, Maupin-Furlow J, Pohlschroder M, Daniels C, Pfeiffer F, Allers T, Eisen JA. 2010. The Complete Genome Sequence of Haloferax volcanii DS2, a Model Archaeon. PLoS ONE 5:e9605.

17. Hawkins M, Malla S, Blythe MJ, Nieduszynski CA, Allers T. 2013. Accelerated growth in the absence of DNA replication origins. Nature 503:544–547.

18. Legerme G, Yang E, Esquivel R, Kiljunen S, Savilahti H, Pohlschroder M. 2016. Screening of a Haloferax volcanii Transposon Library Reveals Novel Motility and Adhesion Mutants. Life 6:41.

19. Nagel C, Machulla A, Zahn S, Soppa J. 2019. Several One-Domain Zinc Finger µ-Proteins of Haloferax Volcanii Are Important for Stress Adaptation, Biofilm Formation, and Swarming. Genes 10:361.

20. Schulze S, Pfeiffer F, Garcia BA, Pohlschroder M. 2021. Comprehensive glycoproteomics shines new light on the complexity and extent of glycosylation in archaea. PLOS Biol 19:e3001277.

21. Jumper J, Evans R, Pritzel A, Green T, Figurnov M, Ronneberger O, Tunyasuvunakool K, Bates R, Žídek A, Potapenko A, Bridgland A, Meyer C, Kohl SAA, Ballard AJ, Cowie A, Romera-Paredes B, Nikolov S, Jain R, Adler J, Back T, Petersen S, Reiman D, Clancy E, Zielinski M, Steinegger M, Pacholska M, Berghammer T, Bodenstein S, Silver D, Vinyals O, Senior AW, Kavukcuoglu K, Kohli P, Hassabis D. 2021. Highly accurate protein structure prediction with AlphaFold. Nature 596:583–589.

22. Holm L. 2022. Dali server: structural unification of protein families. Nucleic Acids Res 50:W210–W215.

23. Garner EC. 2021. Toward a Mechanistic Understanding of Bacterial Rod Shape Formation and Regulation. Annu Rev Cell Dev Biol 37:1–21.

24. Bork P, Sander C, Valencia A. 1992. An ATPase domain common to prokaryotic cell cycle proteins, sugar kinases, actin, and hsp7O heat shock proteins. Proc Natl Acad Sci USA.

25. Cerletti M, Martínez MJ, Giménez MI, Sastre DE, Paggi RA, De Castro RE. 2014. The LonB protease controls membrane lipids composition and is essential for viability in the extremophilic haloarchaeon *H aloferax volcanii*: Characterization of *lonB* mutants in *Haloferax volcanii*. Environ Microbiol 16:1779–1792.
